# OrtSuite – from genomes to prediction of microbial interactions within targeted ecosystem processes

**DOI:** 10.1101/2021.06.04.447094

**Authors:** João Pedro Saraiva, Alexandre Bartholomäus, René Kallies, Marta Gomes, Marcos Bicalho, Carsten Vogt, Antonis Chatzinotas, Peter Stadler, Oscar Dias, Ulisses Nunes da Rocha

## Abstract

The high complexity found in microbial communities makes the identification of microbial interactions challenging. To address this challenge, we present OrtSuite, a flexible workflow to predict putative microbial interactions based on genomic content of microbial communities and targeted to specific ecosystem processes. The pipeline is composed of three user-friendly bash commands. OrtSuite combines ortholog clustering with genome annotation strategies limited to user-defined sets of functions allowing for hypothesis-driven data analysis such as assessing microbial interactions in specific ecosystems. OrtSuite matched, on average, 96 % of experimentally verified KEGG orthologs involved in benzoate degradation in a known group of benzoate degraders. Identification of putative synergistic species interactions was evaluated using the sequenced genomes of an independent study which had previously proposed potential species interactions in benzoate degradation. OrtSuite is an easy to use workflow that allows for rapid functional annotation based on a user curated database and can easily be extended to ecosystem processes where connections between genes and reactions are known. OrtSuite is an open-source software available at https://github.com/mdsufz/OrtSuite.

## Introduction

In environments where microorganisms play a key role, the microbial community functional potential encompasses the building blocks for all possible interspecies interactions (Maestre *et al*, 2012; Mulder *et al*, 2001). For example, in environments rich in methane, microbial communities are dominated by species with genes encoding proteins involved in methanogenesis (Lyu *et al*, 2018). Soil microbes, especially those in the rhizosphere are genetically adapted to support plants in the resistance against pathogens and tolerance to stress (Mendes *et al*, 2018). In this context, natural ecosystems are populated by an enormous number of microbes (Locey & Lennon, 2016). For example, soil environments can contain more than 10^10^ organisms per gram of soil which are distributed in a heterogeneous way making a global search for interspecies interactions unfeasible (Raynaud & Nunan, 2014). The exponential increase in high-throughput sequencing data and the development of computational sciences and bioinformatics pipelines has advanced our understanding of microbial community composition and distribution in complex ecosystems (Roh *et al*, 2010). This knowledge increased our ability to reconstruct and functionally characterize genomes in complex communities, for example by the recovery of metagenome-assembled genomes (MAGs) (Parks *et al*, 2017; Pasolli *et al*, 2019; Tully *et al*, 2018). While several tools have been developed to improve the reconstruction of MAGs, the same cannot be said for predicting interspecies interactions (Morin *et al*, 2018). Studies by Parks (Parks *et al*, 2017) and Tully (Tully *et al*, 2018), while advancing the reconstruction of MAGs, did not perform any functional characterization or prediction of interspecies interactions. Pasolli and collaborators (Pasolli *et al*, 2019) performed functional annotation of representative species in their study by employing several tools such as EggNOG (Huerta-Cepas *et al*, 2017), KEGG (Kanehisa *et al*, 2004) and DIAMOND (Buchfink *et al*, 2015). However, the sheer number of representative genomes (4930) and the lack of focus on specific ecosystem processes makes predicting interspecies interactions a challenge. Furthermore, the challenge of predicting interspecies interactions increases due to the multitude of potential interactions not only between species in microbial communities but also between microbes and their hosts (e.g., plants, animals and microeukaryotes) (Slade *et al*, 2017). An integrated pipeline for annotation and visualization of metagenomes (MetaErg) developed by Dong and Strous (Dong & Strous, 2019) attempt to address some of the challenges in metagenome annotation such as the inference of biological functions and integration of expression data. MetaErg performs comprehensive annotation and visualization of MAGs by integrating data from multiple sources such as Pfam (Mistry *et al*, 2021), KEGG (Kanehisa *et al*, 2004) and FOAM (Prestat *et al*, 2014). However, MetaErg’s full genome annotation requires elevated processing times and computational resources due to its untargeted approach. Furthermore, there is a lack of a user-friendly tool to explore the results tables and graphs to extract pathway specific information tied to each MAG and thus infer potential species interactions based on their functional profiles.

Genome-based modelling approaches have routinely been used to study single organisms as well as microbial communities (Gottstein *et al*, 2016). For example, constraint-based models are highly employed in the study and prediction of metabolic networks (Heirendt *et al*, 2019). These models are generated upon the premise that any given function is feasible as long as the protein-encoding gene is present. Although species may lack the genetic potential to perform all functions necessary to survive in a given ecosystem, in nature microbes do not exist in isolation and may benefit from their interaction with other species. By assessing the genomic content of individual species, we are able to identify groups of microbes whose combined content may account for complete ecosystem functioning. However, generating full genome metabolic networks for each species in a microbial community is time consuming and requires information not easily obtained for each community member such as biomass composition and nutritional requirements.

In order to decrease complexity and facilitate analysis, the search of interactions can be limited to groups of organisms (e.g. microbe-microbe or host-microbe) or specific ecosystem processes (e.g. nitrification or deadwood decomposition). A network-based tool for predicting metabolic capacities of microbial communities and interspecies interactions (NetMet) was recently developed by Tal et al., (Tal *et al*, 2020). The tool only requires a list of species-specific enzyme identifiers and a list of compounds required for a given environment. However, besides the necessity of previous annotation of genomes, NetMet does not consider the rules that govern each reaction (e.g. protein complexes). Accurate annotation of gene function from sequencing data is essential to predict, ecosystem processes potentially performed by microbial communities, particularly in cases where an ecosystem process is performed by the synergy of two or more species. Simple methods for the annotation of genomes rely, for instance, on the search for homologous sequences. Computational tools such as BLAST (Altschul *et al*, 1990) and DIAMOND (Buchfink *et al*, 2015) allow the comparison of nucleotide or protein sequences to those present in databases. These approaches allow inferring the function of uncharacterized sequences from their homologous pairs whose function is already known. The degree of confidence in the assignment of biological function is increased if this has been validated by, for example, experimental data. Approaches based on orthology are increasingly used for genome-wide functional annotation (Huerta-Cepas *et al*, 2017). Orthologs are homologous sequences that descend from the same ancestor separated after a speciation event retaining the same function (Koonin, 2005). OrthoMCL (Li *et al*, 2003), CD-HIT (Li & Godzik, 2006) and OrthoFinder (Emms & Kelly, 2015, 2019) are just a few tools that identify homologous relationships between sequences using orthology. OrthoFinder has been shown to be more accurate than several other orthogroup inference methods since it considers gene length in the detection of ortholog groups by introducing a score transformation step (Emms & Kelly, 2015). However, OrthoFinder, due to its all-versus-all sequence alignment approach, requires intensive computational resources resulting in long running times when using large data sets for clustering. Because of the enormous number of potential combination, limiting the scope of research to specific ecosystem processes may reduce the computational and resource costs associated with the integration of ortholog clustering tools and functional annotation strategies.

In this study, we developed OrtSuite; a workflow that can: (i) perform accurate ortholog based functional annotation, (ii) reveal putative microbial synergistic interactions, and (iii) digest and present results for pathway and community driven biological questions. These different features can be achieved with the use of three bash commands in a reasonable computational time. This research question / hypothesis targeted approach integrates a user-defined database – Ortholog-Reaction Association database (ORAdb) – with up-to-date ortholog clustering tools. OrtSuite allows the search for putative microbial interactions by calculating the combined genomic potential of individual species in specific ecosystem processes. OrtSuite also provides a visual representation of the species genetic potential mapped to each of the reactions defined by the user. We evaluate this workflow using a clearly defined set of reactions involved in the well-described benzoate-to-Acetyl-CoA (BTA) conversion. Further, we used this workflow to functionally characterize a set of known benzoate degraders. OrtSuite’s ability to identify putative interspecies interactions was evaluated on species whose potential interactions have been previously predicted under controlled conditions (Fetzer *et al*, 2015).

## Results

### Ortsuite is a flexible and user-friendly pipeline

One of the motivations to develop Ortsuite was to facilitate the targeted analysis of the genomic potential of microbial communities including the prediction of putative synergistic interspecies interactions. To achieve this, OrtSuite was developed to integrate ortholog clustering tools (Emms & Kelly, 2019) with sequence alignment programs (Buchfink *et al*, 2015). To increase user-friendliness, three scripts were created that, collectively, perform all five tasks associated with OrtSuite: (1) download of sequences to populate ORAdb, (2) generation of Gene-Protein-Reaction (GPR) rules, (3) clustering of orthologs, (4) targeted functional annotation and (5) prediction of putative synergistic interspecies interactions (Figure 1). Additional control is also given to the user such as establishing thresholds in the minimum e-values (during sequence alignment of sequences in ortholog clusters to ORAdb). Other constraints include restricting the number of putative microbial interactions based on the presence of transporters and subsets of reactions to be performed by individual species (Supplementary data – Table S1). Since data in public repositories is frequently being added or updated and to include personal knowledge the user can manually curate the files in the ORAdb and GPR rules, with the latter being strongly advised.

**Figure 1.**
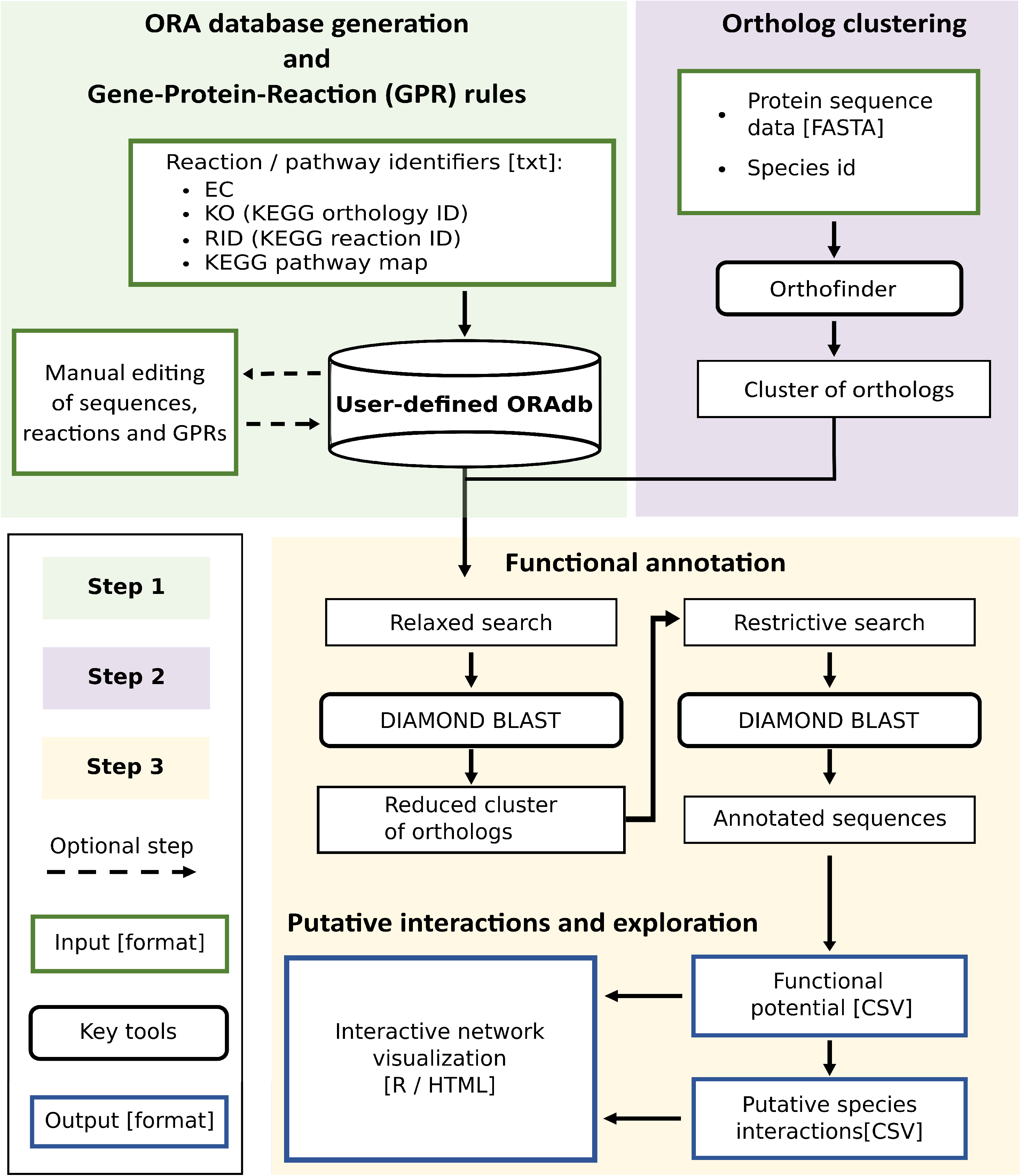
OrtSuite workflow. OrtSuite takes a text file containing a list of identifiers for each reaction in the pathway of interest supplied by the user to retrieve all protein sequences from KEGG Orthology and are stored in ORAdb. Subsequently the same list of identifiers is used to obtain the Gene-Protein-Reaction (GPR) rules from KEGG Modules (Step 1). Protein sequence from samples supplied by the user are clustered using OrthoFinder (Step 2). In step 3, the tasks of functional annotation, identification of putative synergistic species interactions and graphical visualization of the network are performed. Functional annotation consists of a two-stage process (relaxed and restrictive search). Relaxed search performs sequence alignments between 50% of randomly selected sequences from each generated cluster. Clusters whose representative sequences share a minimum E-value of 0.001 to sequences in the reaction set(s) in ORAdb transition to the restrictive search. Here, all sequences from the cluster are aligned to all sequences in the corresponding reaction set(s) to which they had a hit (default E-value = 1e^-9^). Next, the annotated sequences are further filtered to those with a bit score greater than 50 and are used to identify putative microbial interactions based on their functional potential. Constraints can also be added to reduce the search space of microbial interactions (e.g. subsets of reactions required to be performed by single species, transport-related reactions). Additionally, an interactive network visualization of the results is produced and accessed via a HTML file.

A git repository for OrtSuite (https://github.com/mdsufz/OrtSuite) was also generated. This repository provides users with an easy-to-follow detailed guide covering installation to the running of the three scripts and generated outputs.

### Computing time of OrtSuite stages

The runtime of each OrtSuite step was evaluated on a set of genomes whose genomic potential in the conversion of benzoate to acetyl-CoA was known (Table 1). The same set of genomes is used in the OrtSuite’s GitHub tutorial page (https://github.com/mdsufz/OrtSuite/blob/master/OrtSuite_tutorial.md) with a total of 75.5 Megabytes of data. OrtSuite was used to analyze this data on a laptop with 4 cores and 16 Gigabytes of RAM. All OrtSuite steps were run on default settings, and the total runtime of each step was recorded (Table 2). The total workflow was completed in 3 h 50 min and the longest single step runtime consisted of 2 h and 47 min which involved the construction of the ORAdb. The user does have the option to modify the number of cores used during functional annotation which should further decrease run times.

**Table 1.** Species names, strain and abbreviation codes of the genomes used to validate OrtSuite (Supplementary data - Test_genome_set). The genomic potential, based on KEGG database, to completely encode all proteins involved in a BTA pathway is identified in the column “BTA pathway” (P1 – Anaerobic conversion of benzoate to acetyl-CoA 1; P2 – Anaerobic conversion of benzoate to acetyl-CoA 2; P3 – Aerobic conversion of benzoate to acetyl-CoA). * indicates no literature was found connecting benzoate degradation and the respective species.

| Name and strain | Abbreviation code | KEGG id | BTA pathway | Accession number | Ref. |
| --- | --- | --- | --- | --- | --- |
| *Acinetobacter defluvii* WCHA30 | adv | T05474 | P3 | CP029389-CP029397 | (Hu *et al*, 2017) |
| *Arabidopsis thaliana* | ath | T00041 | - | GCF_000001735 | * |
| *Azoarcus sp.* KH32C | aza | T02502 | P2 | AP012304, AP012305 | (Junghare *et al*, 2015) |
| *Azoarcus sp.* DN11 | azd | T05691 | P2 | CP021731 | (Devanadera *et al*, 2019) |
| *Azoarcus sp.* CIB | azi | T04019 | P2 | CP011072 | (Valderrama *et al*, 2012) |
| *Burkholderia cepacia* DDS 7H-2 | bced | T03302 | P3 | CP007785-CP007787 | (Jenul *et al*, 2018) |
| *Burkholderia vietnamiensis* G4 | bvi | T00493 | P3 | CP000614-CP000621 | (O’Sullivan *et al*, 2007) |
| *Cycloclasticus sp.* P1 | cyq | T02265 | P3 | CP003230 | (Wang *et al*, 2008) |
| *Cycloclasticus zancles* 78-ME | cza | T02780 | P3 | CP005996 | (Messina *et al*, 2016) |
| *Desulfosporosinus orientis* DSM 765 | dor | T01675 | - | CP003108 | (Robertson *et al*, 2000) |
| *Aromatoleum aromaticum* EbN1 | eba | T00222 | P2 | CR555306-CR5553068 | (Rabus *et al*, 2016) |
| *Latimeria chalumnae* (coelacanth) | lcm | T02913 | - | GCF_000225785 | * |
| *Magnetospirillum sp.* XM-1 | magx | T04231 | P2 | LN997848-LN997849 | (Meyer-Cifuentes *et al*, 2017) |
| *Paraburkholderia aromaticivorans* BN5 | parb | T05169 | P3 | CP022989-CP022996 | (Lee *et al*, 2019) |
| *Rhodococcus ruber* P14 | rrz | T05142 | P3 | CP024315 | (Peng *et al*, 2018) |
| *Sulfuritalea hydrogenivorans* sk43H | shd | T03591 | P2 | AP012547 | (Sperfeld *et al*, 2019) |
| *Staphylococcus sciuri* FDAARGOS 285 | sscu | T05176 | - | CP022046-CP022047 | (Mrozik & Labuzek, 2002) |
| *Thauera sp.* MZ1T | tmz | T00804 | P2, P3 | CP001281-CP001282 | (Suvorova & Gelfand, 2019) |

**Table 2.**
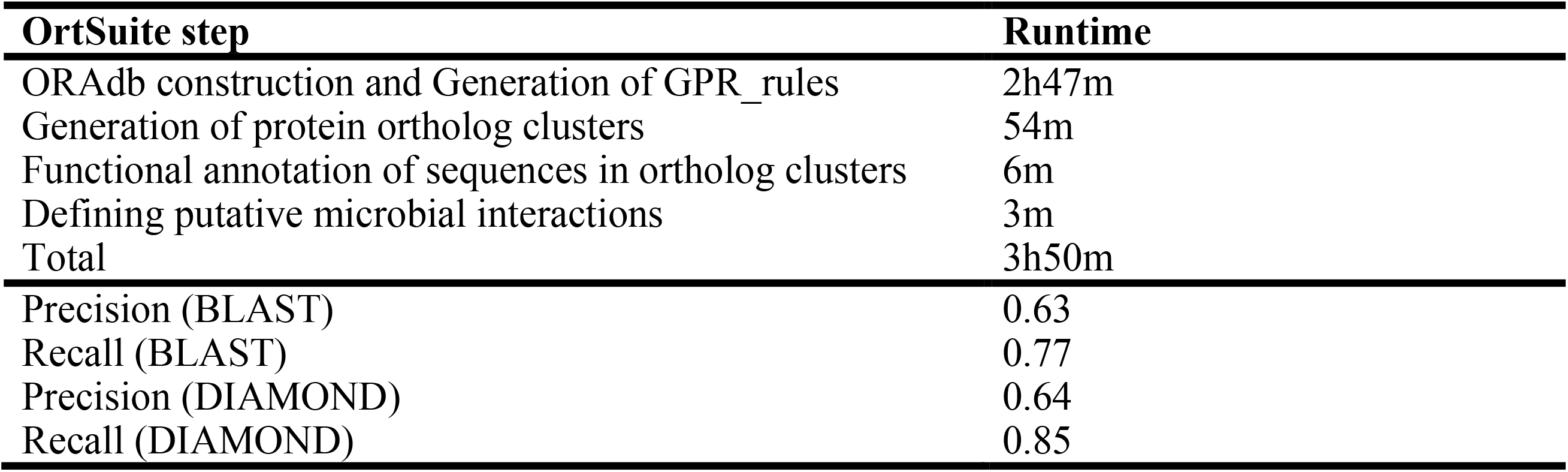
OrtSuite workflow runtime and clustering performance. The total runtime of each OrtSuite step when analyzing the genomic potential of species in Test_genome_set dataset in three pathways (P1, P2 and P3) for the conversion of benzoate to acetyl-CoA (BTA). Steps were performed with default parameters on a laptop with 4 cores and 16 GB of RAM. Pair-wise precision and recall results of OrthoFinder using BLAST and DIAMOND as an alignment search tool. Clustering was performed on the Test_genome_set dataset plus the mutated genomes.

| **OrtSuite step** | **Runtime** |
| --- | --- |
| ORAdb construction and Generation of GPR_rules | 2h47m |
| Generation of protein ortholog clusters | 54m |
| Functional annotation of sequences in ortholog clusters | 6m |
| Defining putative microbial interactions | 3m |
| Total | 3h50m |
| Precision (BLAST) | 0.63 |
| Recall (BLAST) | 0.77 |
| Precision (DIAMOND) | 0.64 |
| Recall (DIAMOND) | 0.85 |

### Higher recall rates during clustering of orthologs with DIAMOND

We performed an evaluation of the effects of point mutations during clustering of orthologs using OrthoFinder (Emms & Kelly, 2019). OrthoFinder allows users to choose between DIAMOND (Buchfink *et al*, 2015) and BLAST (Altschul *et al*, 1990) as sequence aligners. To test which sequence aligner yielded the best results we performed ortholog clustering of a dataset consisting of the original target genomes as well as a set of artificially mutated genomes (Supplementary data - Test_genome_set) using both aligners. The results showed 0.01 difference between OrthoFinder and DIAMOND precision (Table 2). However, DIAMOND showed a 9.5% higher recall than that observed for OrthoFinder what suggests DIAMOND may have higher sensitivity in the clustering of sequences with the same function. All artificially mutated sequences (even those with mutation rates of 25%) were clustered together with their non-mutated ortholog. In parallel, we also performed sequence alignment using NCBI’s BLASTp (Madden, 2003) between the protein sequences of the DNA-mutated and un-mutated genes. E-values of sequence alignments in all species ranged from 0 to 5e^-180^ and percentage of identity from 61.32 to 98.84% (Supplementary data – Table S2). For validation of the OrtSuite workflow, clustering of protein orthologs was repeated using only the original unmutated 18 genomes and the default aligner (DIAMOND). A complete overview of the results generated during the clustering of orthologs (e.g. number of genes in ortholog clusters, number of unassigned genes and number of ortholog clusters) was also obtained (Supplementary data - Table S3).

### High rate of KEGG annotations predicted by OrtSuite

The third step of OrtSuite consists of performing cluster annotation in a two-stage process. In the first, only 50% of sequences are used in the alignment to the sequences ORAdb. Those with a minimum e-value proceed to the second stage where all sequences contained in this cluster will be aligned. At the end, annotation of clusters will take into consideration additional parameters such as bit scores. To evaluate the thresholds used in the annotation of ortholog clusters we used one relaxed (0.001) and four restrictive (1e^-4^, 1e^-6^, 1e^-9^ and 1e^-16^) e-value cutoffs. An overview of the results (e.g. number of clusters containing orthologs from ORAdb, number of ortholog clusters with annotated sequences) is shown in (Supplementary data – Table S4). The performance of OrtSuite in the functional annotation of the genomes in the *Test_genome_set* is shown in (Supplementary data – Table S5). On average, 96% of the annotations assigned by KEGG were also identified by OrtSuite. The complete list of results of functional annotation using the different e-value cutoffs are available in the Supplementary data - Table S6, Table S7, Table S8 and Table S9. Similarly, the mapping of species with the genetic potential for each reaction (considering the GPR rules) using the different e-value cutoffs can be found in the Supplementary data – Table S10, Table S11, Table S12 and Table S13. In terms of annotation, no striking difference was observed between the four different e-value cutoffs used during the restrictive search stage. However, the largest decrease in the number of ortholog clusters that transits from the relaxed search to the restrictive occurs when using an e-value cutoff of 1e^-16^ (Supplementary data – Table S4). The difference in computing time between lower and higher e-value thresholds was negligible (< 2 min). Other annotation tools, such as NCBI’s BLAST tool (Altschul *et al*, 1990), BlastKOALA (Kanehisa *et al*, 2016) and Prokka (Seemann, 2014), can annotate full genomes, the latter at a relatively fast pace. On average, full genome annotation of our genomes in the *Test_genome_set* dataset using Prokka required 12 mins on a customary laptop with 16 Gigabytes of RAM and four CPUs to complete. BlastKOALA required approximately 3 hours to annotate a single genome.

### Identifying genetic potential to perform a pathway

To test OrtSuite’s ability to identify species with the genetic potential to perform a pathway individually we defined sets of reactions that are used in three alternative pathways for the conversion of benzoate to acetyl-CoA (Supplementary data – Table S14). Next, we compared the results to the species’ known genomic content in each alternative pathway (Supplementary data – Table S15). OrtSuite matched KEGG’s predictions in species’ ability to perform each alternative benzoate degradation pathway in all but two species - *Azoarcus* sp. DN11 and *Thauera* sp. MZ1T. Furthermore, OrtSuite identified five species capable of performing conversion pathways not contemplated in KEGG. *Azoarcus sp*. KH32C, *Aromatoleum aromaticum* EbN1, *Magnetospirillum sp*. XM-1 and *Sulfuritalea hydrogenivorans* sk43H have the genetic potential to perform both pathways involving the anaerobic conversion of benzoate to acetyl-CoA while *Azoarcus sp*. CIB has to genetic potential to perform all alternative pathways (except when using an e-value cutoff of 1e^-16^). No genes in *Thauera sp*. MZ1T involved in the conversion of crotonyl-CoA to 3-Hydroxybutanoyl-CoA (R03026) were identified by OrtSuite which impedes the anaerobic conversion of benzoate to acetyl-CoA. The default e-value for the restrictive search was set to 1e^-9^ since OrtSuite’s performance did not change significantly between all tested e-value cutoffs but showed a greater drop in the number of consistent orthogroups (i.e. clusters of orthologs whose sequences are all annotated with the same function) from 1e^-9^ to 1e^-16^.

### Using OrtSuite to predict interspecies interactions

In this study, we tested the ability of OrtSuite in identifying interspecies interactions involved in the conversion of benzoate to acetyl-CoA where experimental data were available. Prediction of synergistic interspecies interactions was assessed on a set of sequenced isolates (Supplementary data - Fetzer_genome_set.zip): Monocultures of these isolates and randomly assembled communities of one to 12 species including these benzoate-degraders and additional species incapable of directly using benzoate as a carbon source were analyzed previously (Fetzer *et al*, 2015) under three different environmental conditions (low substrate concentration:1g/L benzoate, high substrate concentration: 6g/L benzoate and high substrate concentration + additional osmotic stress: 6g/L benzoate supplemented with 15g /L of NaCl). In that study, Fetzer et al investigated if the presence or absence of a particular species positively or negatively affected biomass production. Since under specific conditions the presence of a degrader alone was not sufficient for community biomass production, they further analyzed if potential species interactions could be of relevance. Briefly, they defined for all environmental conditions minimal communities, which showed community growth without the need to include other species and identified whether the presence of a single species alone or potential interaction between the specific species (and thus potential partners) present in these minimal communities stimulated biomass production (Fetzer *et al*, 2015). Using OrtSuite, we aimed to identify which potential species interactions predicted by Fetzer and collaborators could be a result of their combined genetic potential.

Our dataset contained 69,193 protein sequences distributed across the 12 species resulting in a total of 59 Megabytes of data. More than 84% of all genes were placed in 9,533 ortholog clusters. In addition, 541 clusters were composed of sequences obtained from all 12 species (Supplementary data - Table S16). OrtSuite’s annotation stage resulted in 326 ortholog clusters with annotated sequences from ORAdb (Supplementary data - Table S17). The mapping of KOs to each species in the *Fetzer_genome_set* is available as supplementary data (Table S18). The genomic potential of each species for aerobic and anaerobic benzoate metabolizing pathways is shown in Figure 2. The complete mapping of reactions to each species is available in the supplementary data (Table S19). Based on the 326 ortholog clusters and the Gene-Protein-Reaction (GPR) rules (Supplementary data - Table S20), five species (*Cupriavidus necator* JMP134, *Pseudomonas putida* ATCC17514, *Rhodococcus sp*. Isolate UFZ (Umweltforschung Zentrum), *Rhodococcus ruber* BU3 and *Sphingobium yanoikuyae* DSM6900) contained all protein-encoding genes required to perform aerobic conversion of benzoate to acetyl-CoA. In the Fetzer study, *Rhodococcus sp*. Isolate UFZ and *S. yanoikuyae* did not show growth in a medium containing benzoate. The incomplete functional potential of *C. testosteroni* ATCC 17713 and *P. putida* ATCC17514 to perform aerobic conversion of benzoate to acetyl-CoA is at odds with their reported growth as monocultures in the presence of benzoate as shown in the Fetzer study. The number of species with the genetic potential for each reaction involved in the aerobic benzoate degradation pathway (P3) is shown in (Supplementary data - Table S21). Usually, all species with the complete genomic potential to perform a complete pathway are excluded when calculating interspecies interactions since they do not require the presence of others. However, species identified by OrtSuite with the complete functional potential to perform each defined pathway were also included to compare to the results in the Fetzer study presented above. A total of 2382 combinations of species interactions were obtained whose combined genetic potential covered all reactions. The complete list of potentially interacting species is available in the supplementary data (Table S22).

**Figure 2.**
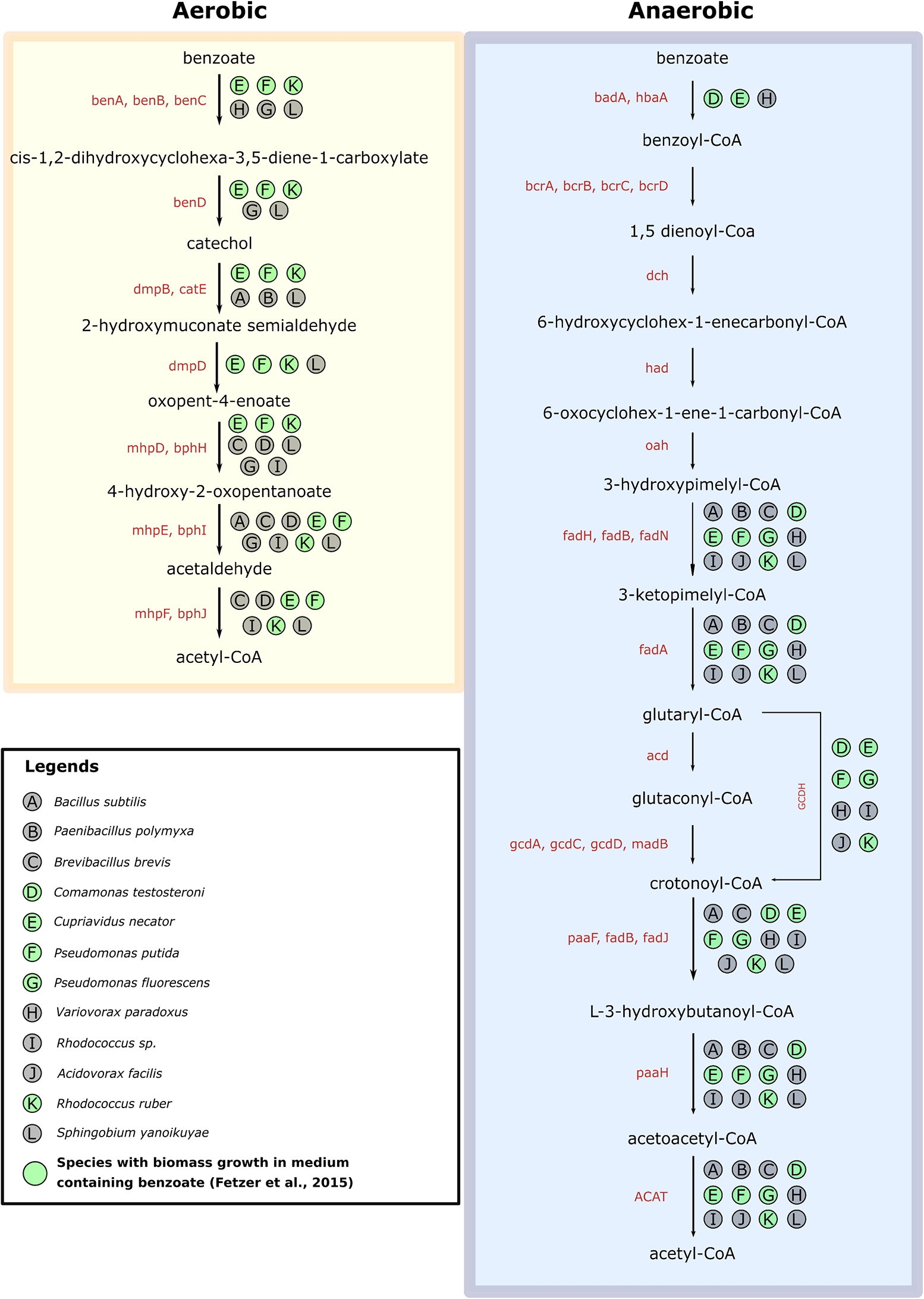
Mapping of the genomic potential of each species from the Fetzer_genome_set dataset to each reaction in aerobic (yellow) and anaerobic (blue) benzoate-to-acetyl-CoA conversion pathways. Circles highlighted in green represent species that showed biomass growth in medium containing benzoate in the Fetzer study.

**Figure 3.**
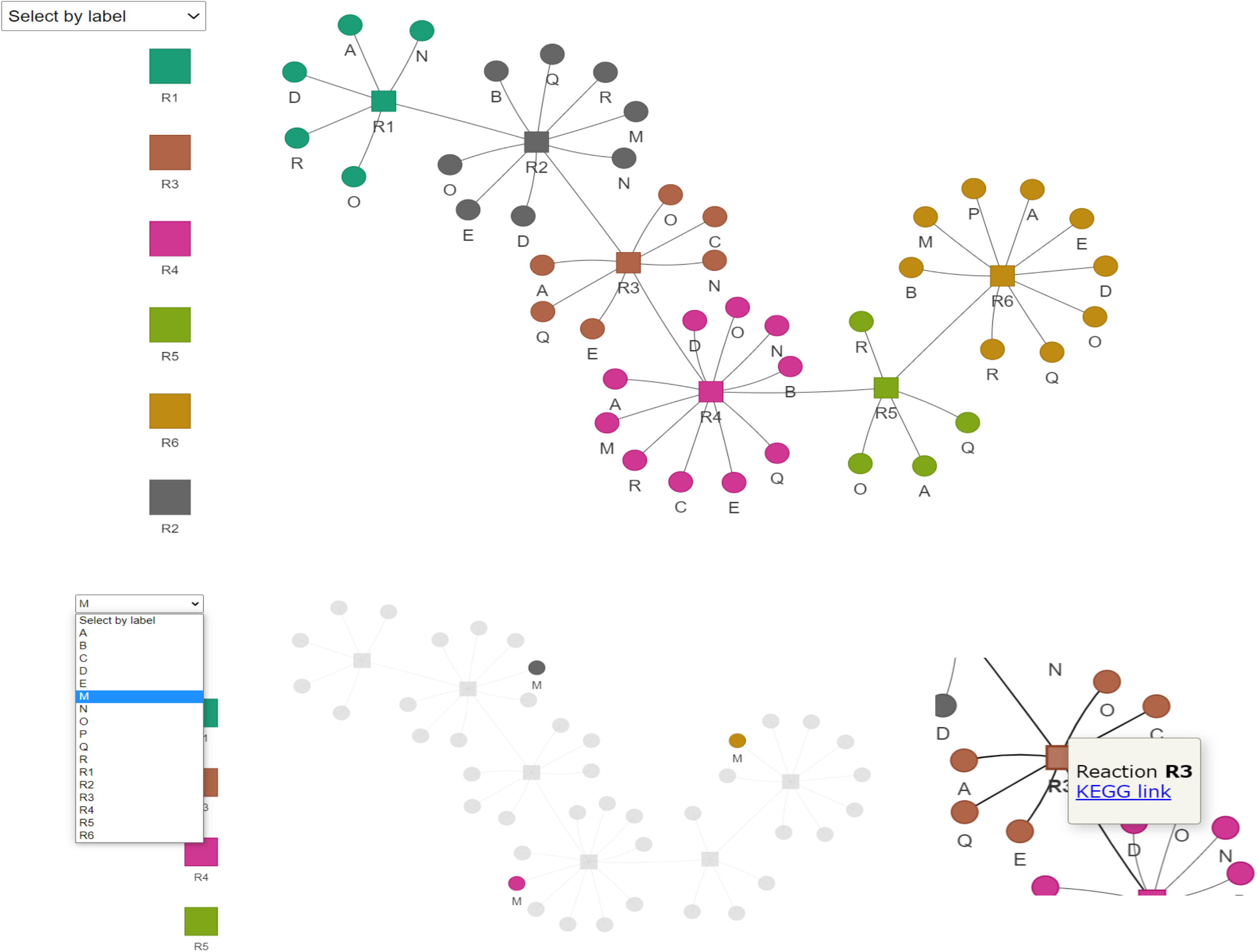
Example of the interactive network visualization included on OrtSuite results. (A) The complete network with species colored by reaction (B) Species can be highlighted for simple identification (C). Tooltips on reaction link out the KEGG if the reaction identifier is given.

In the anaerobic degradation pathways (P1 and P2) no species presented the genomic content to encode proteins involved in the conversion of benzoyl-CoA to Cyclohexa-1,5-diene-1-carboxyl-CoA (R02451) (Supplementary data - Table S23). This reaction requires the presence of a protein complex either composed of four subunits (K04112, K04113, K04114, K04115) or composed of two subunits (K19515, K19516). No species was annotated with all subunits in either protein complex. Therefore, no species interactions were identified that would allow the complete anaerobic conversion of benzoate to acetyl-CoA. In the low substrate environment, OrtSuite identified 826 of 830 (99.5%) species combinations showing growth. In the high substrate environment, OrtSuite predicted 644 of 646 (99.7%). In the high substrate+salt stress environment, OrtSuite predicted all 271 (100%) combinations of species exhibiting growth (Supplementary data - Table S24).

## Discussion

We designed OrtSuite to allow hypothesis-driven exploration of microbial interactions in a user-friendly manner. This was achieved by integrating up-to-date clustering tools with faster sequence alignment methods and limiting the scope to user-defined ecosystem processes or metabolic functions. Using only three bash commands required to run the complete workflow, OrtSuite is a user-friendly tool capable of running in a customary computer (four cores and 16GB of RAM) with even faster runtimes when using high performance computing.

The clustering of orthologs by OrthoFinder using DIAMOND (Buchfink *et al*, 2015) showed higher sensitivity and lower runtime compared to BLAST (Altschul *et al*, 1990) which has also been shown by Hernández-Salmerón and Moreno-Hagelsieb (Hernández-Salmerón & Moreno-Hagelsieb, 2020). Furthermore, low e-values and medium to high identity percentages in the sequence alignments between mutated and original genes indicates that the mutated genes still share enough sequence similarity to the original protein sequence. These results suggest that mutation rates of up to 25% of single DNA base pairs will not have an observable effect on the clustering of orthologs. OrthoFinder’s algorithm removes the gene length bias from the sequence alignment process, which may also explain why mutated genes were clustered with the original. Although it has been suggested that most genetic variations are neutral, changes in single base pairs can have a drastic effect on protein function (e.g. depending on the location of the mutation) (Ng & Henikoff, 2006). To this purpose, experimental functional studies can be used to validate previously unannotated orthologs. Furthermore, this study case does not consider the distribution of mutations across species and gene families which can also have different effects on the clustering of orthologs (Khanal *et al*, 2015). Therefore, future studies increasing the rates of DNA base pair substitutions and other types of mutations as well as experiments targeting protein function in ortholog clusters are needed.

Next, we aimed to improve and facilitate functional annotation and prediction of synergistic microbial interactions. Exploring the great amount of data generated from full genome annotation of individual species from complex microbial communities is a daunting task. This is evident in a study by Singleton and collaborators (Singleton *et al*, 2021) where the connection between structure and function required the analysis of metagenomics data, 16S and molecular techniques such as fluorescent in situ hybridization and Raman spectroscopy. Looking solely to functional annotation, two challenges, among others, arise: First, performing all vs all sequence alignments in complex communities is resource consuming (time and computational power) and second, manual inspection of each annotated genome for target genes or pathways is required. Identifying interspecies interactions based on the microbe’s complete genomic potential is also challenging. For example, network approaches are increasingly employed in ecology but selection of the most appropriate approach is not always straightforward and easy to implement (Delmas *et al*, 2019). OrtSuite overcomes these challenges by first performing cluster annotation in a two-stage process and limited to user-defined set of functions of interest which decreases the number of sequence alignments to be performed. The user-defined database coupled with the scripts for automated identification of interspecies interactions contained in OrtSuite decreases the time required to generate the data and facilitates its interpretation by the user. Additionally, OrtSuite generates a graphical representation of the network further facilitating analysis of the whole microbial community (https://github.com/mdsufz/OrtSuite/blob/master/network_example.png).

OrtSuite not only confirmed all but two of KEGG’s predictions in species’ ability to perform each alternative benzoate degradation pathway used in this study but also identified five species capable of performing conversion pathways not contemplated in KEGG. On average, an additional 18.3 KO identifiers were mapped to genes not previously annotated in our species. The use of e-value and bit score as the filtering criteria rather than sequence identity, employed by KEGG, may explain the increase in functionally annotated genes. For example, the alignment of a sequence of *A. defluvii* (adv: AWL30228.1) to the sequences in ORAdb annotated as K04105 (conversion of benzoate to benzoyl-CoA) showed high bit-scores (200.7) and low e-values (2e^-54^) but the identity percentage did not exceed 28.6%.The use of e-values and bit scores to infer function has been nicely reviewed by Pearson (Pearson, 2013) who suggests that e-values and bit scores are more sensitive and reliable than identity percentages in finding homology since they take into account evolutionary distance of aligned sequences, the sequence lengths and the scoring matrix.

To test the prediction of putative synergistic microbial interactions we used data from an independent study performed by Fetzer and collaborators (Fetzer *et al*, 2015); hereafter Fetzer study. In the Fetzer study five species showed biomass growth (estimated by optical density at 590nm wave lenght) in medium containing benzoate. We evaluated whether these species possessed the complete genomic content to encode all proteins required for each benzoate to acetyl-CoA conversion pathway. The remaining seven species were not able to grow as monocultures in media with benzoate as sole carbon source. Therefore, we evaluated whether the lack of growth was confirmed by lack of essential protein-encoding genes involved in conversion of benzoate to acetyl-CoA. Fetzer study also showed that, under specific nutrient and stress conditions, total biomass production was influenced by the presence of non-degrading species. Thus, we evaluated whether putative species interactions identified by OrtSuite fit the results obtained by in the Fetzer study. OrtSuite confirmed the functional potential for aerobic conversion of benzoate to acetyl-CoA in three of the five species whose growth in monocultures was observed during their study. In Fetzer’s study, two species, *S. yanoikuyae* (accession number GCA_903797735.1) and *Rhodococcus sp*. (accession number GCA_903819475.1), were not able to grow as monoculture in the presence of benzoate. However, OrtSuite predicted that both possessed the functional potential to aerobically convert benzoate to acetyl-CoA. In their study, in a medium containing 1g/L of benzoate, growth was considered when optical densities (OD) were above 0.094. The OD measured for *S. yanoikuyae* was 0.0916. The annotation of genes with the ability to perform the complete aerobic conversion of benzoate to acetyl-CoA combined with the small difference in OD to the minimum threshold suggests that *S. yanoikuyae* indeed can grow on low benzoate containing medium but at perhaps at lower growth rates. In the case of *Rhodococcus sp*. Isolate UFZ, the OD was never measured above 0.022 which, again, might indicate slow growing species. Another possible explanation is that although these two species possess the genes necessary for aerobic benzoate degradation they are not active. In Fetzer’s study, the observed growth of *Comamonas testosteroni* ATCC11996 and *Pseudomonas fluorescens* DSM6290 in the low benzoate environment was not confirmed by OrtSuite. To note, benzoate conversion intermediates were not determined in the Fetzer experiment. Hence, it is possible that these two species utilize reactions or pathways that were not included in the benzoate degradation pathways used in our study. Despite the presence of benzoate degraders, another possible explanation as to the unobserved growth in Fetzer’s study for certain experimental conditions is the lack of tolerance of these species to high benzoate concentrations. For example, *C. necator* growth was shown to be stimulated at low benzoic acid concentrations but inhibited at high concentrations (Wang *et al*, 2014). In addition, the set of genes used in our study did not consider the presence of stress related factors. To assess these effects, stress-resistance associated genes and reactions such as those involved in medium acidification (Kitko *et al*, 2009) could be added as constraints. Similar results were obtained when using a high substrate+salt stress medium. Under these conditions, presence of benzoate degraders alone was not sufficient to achieve growth of species combinations. Benzoate degradation has been shown to decrease in hyperosmotic environments (Bazire *et al*, 2007) therefore, additional constraints such as genes that confer resistance to environmental stressors or adverse conditions sodium chloride (NaCl) could be included during the identification of interspecies interactions under different or changing environmental conditions.

No single species or combination of species possessed the complete genomic potential to anaerobically convert benzoate to acetyl-CoA via the two proposed pathways (P1 and P2). Since all growth experiments were conducted in aerobic conditions, it is possible that the species in question are only capable of using benzoate as a carbon source in aerobic environments. To fully explore all the species potential to convert benzoate, additional degradation pathways could be could be checked in the future using a multi-omics approach. Furthermore, the only constraints added were related to the reactions that composed each pathway. Additional constraints can be included in future studies, such as potential mandatory transport-associated reactions, to increase confidence in the proposed interspecies interactions. OrtSuite confirmed that most interspecies interactions (> 99%) identified by Fetzer and collaborators were possible due to their combined metabolic potential to aerobically degrade benzoate to acetyl-CoA but not under anoxic conditions.

In this study, we ran OrtSuite on a dataset comprised of 18 genomes (Table 1). To determine if this range would be within the number of genomes in regular microbiome studies we calculated the average number of MAGs from different studies focusing on their recovery. A study performed by Parks and collaborators (Parks *et al*, 2017) analyzed sequencing data from 149 projects. Most projects (91%) consisted of less than 20 samples. On average, they recovered 5.3 metagenome-assembled genomes (MAGs) per metagenome. Work performed by Pasolli and collaborators (Pasolli *et al*, 2019) on microbial diversity in the human microbiome recovered, on average, 16 MAGs per metagenomic library. From the 46 studies used in their work, 30 consisted of less than 200 samples. Another study by Tully and collaborators focusing on marine environments (Tully *et al*, 2018) recovered 2631 MAGs from 234 samples (average of 11 MAGs per sample). Our analysis demonstrates that the average number of MAGs recovered from a metagenome currently range from five to 16. Therefore, performing targeted functional annotation and interspecies interactions predictions using OrtSuite in average sized metagenome samples is still feasible using a customary laptop.

In summary, OrtSuite allows hypothesis-driven exploration of potential interactions between microbial genomes by limiting the search universe to a user-defined set of ecosystem processes. This is achieved by rapidly assessing the genetic potential of a microbial community for a given set of reactions considering the relationships between genes and proteins. The two-step annotation of clusters of orthologs with a personalized ORAdb decreases the overall number of sequence alignments that need to be computed. User-specified constraints, such as the presence of transporter genes, further reduces the search space for putative microbial interactions. Users have substantial control over several steps of OrtSuite: from manual curation of ORAdb, custom sequence similarity cutoffs to the addition of constraints for inference of putative microbial interactions. The reduction of the search space of synergistic interactions by OrtSuite will also allow more comprehensive and computationally demanding tasks to be performed such as (Community) Flux Balance Analysis which depend heavily on genome-scale metabolic models (Thommes *et al*, 2019; Ravikrishnan & Raman, 2021). As long as links between genes, proteins and reactions exist, the flexibility and easy usage of OrtSuite allow its application to the study of any given ecosystem process.

## Materials and Methods

### OrtSuite workflow

The OrtSuite workflow consists of three main steps performed by the use of three bash commands (Figure 1). Briefly, the first step consists in the generation of a user defined ortholog-reaction associated database (ORAdb) and collection of the gene-protein-reaction (GPR) rules. This task takes as input a list of KEGG identifiers which will be used to download all protein sequences associated with a set of reactions/pathway of interest. Next, all gene-protein-rules (GPRs) associated with each reaction will be downloaded from KEGG Modules. In the second step OrtSuite employs OrthoFinder (Emms & Kelly, 2015) to generate ortholog clusters. This step takes as input a folder with the location of the genomic sequences. The third step consists of the functional annotation of species, identification of putative synergistic interspecies interactions and generation of visual representations of the results.

### OrtSuite step 1 (green box, Figure 1) – User defined Ortholog-Reaction Association database (ORAdb) and Gene-Protein-Reaction (GPR) rules file

The ORAdb used for functional annotation consists of sets of protein sequences involved in the enzymatic reactions that compose a pathway/function of interest defined by the user. This database is generated during the execution of the *DB_construction*.*sh* script in OrtSuite requiring only the user to provide:

- a location of the project folder where all results will be stored
- a text file with a list of KEGG identifiers (one identifier per line)
- the full path to the OrtSuite installation folder

The list of identifiers can be KEGG reactions (RID) (e.g. R11353, R02451), enzyme commission (EC) numbers (e.g. 1.3.7.8, 4.1.1.103) or KEGG ortholog identifiers (e.g. K07539, K20941). This file is used by OrtSuite to automatically retrieve the KEGG Ortholog identifiers (KO) (in case the identifiers provided are not KO identifiers) and to download all their associated protein sequences (Kanehisa *et al*, 2004). OrtSuite makes use of the python library *grequests* which allows multiple queries in KEGG subsequently decreasing the time required for retrieving the ortholog associated sequences. The user-defined ORAdb will be composed of KO-specific sequence files in FASTA format associated with all reactions/enzymes of interest. Users also have the opportunity to manually add or edit the sets of reactions and the associated protein sequences in the ORAdb. This feature is of particular importance since many reactions associated with ecosystem processes are constantly being discovered and updated and might not be included in the latest version of KEGG. In addition, during the execution of the D*B_construction*.*sh* OrtSuite performs the automated download of the gene-protein-reaction (GPR) rules from KEGG Modules. This feature is vital since many reactions can be catalyzed by enzymes with a single (i.e., one protein) or multiple subunits (i.e., protein complexes). Despite the automated process, it is strongly advised to manually curate the final table to guarantee accurate results. An example of the final GPR table is shown in the Supplementary data (Table S20).

### OrtSuite step 2 (purple box, Figure 1) - Generation of protein ortholog clusters

The second step of OrtSuite, takes a set of protein sequences and generates clusters of orthologs. This set of protein sequences can originate from single isolates or from the complete set of protein sequences recovered from metagenomes or metagenome-assembled genomes. Indeed, the use of protein sequences from isolates, metagenome-assembled genomes and co-culture experiments will benefit greatly from OrtSuite’s reduction of the universe of potential microbial interactions based on the user defined ORAdb. Orthology considers that phylogenetically distinct species can share functional similarities based on a common ancestor (Gabaldón & Koonin, 2013). Potentially, genes with equal function will be grouped together. To perform this task the OrtSuite pipeline uses OrthoFinder (Emms & Kelly, 2015). Two sequence aligners are available in OrthoFinder – DIAMOND (Buchfink *et al*, 2015) and BLAST (Altschul *et al*, 1990). DIAMOND is used by default due to its improved trade-off between execution time and sensitivity (Emms & Kelly, 2019). This step is performed by running the command *orthofinder* located in the installation folder of OrthoFinder. This command takes as input the full path to the folder containing the protein sequences to be clustered and the full path to the folder where results are to be stored.

### OrtSuite step 3 (yellow box, Figure 1) - Functional annotation of ortholog clusters

The third step of OrtSuite consists in the assignment of functions to protein sequences contained in the ortholog clusters. Functional annotation of these clusters consists of a two-step process termed relaxed and restrictive search, respectively. The goal of the relaxed search is to decrease the number of alignments required to assign functions to sequences in the ortholog clusters. Here, 50% of the total number of sequences from each cluster are randomly selected and aligned to all sequences associated to each reaction present in the ORAdb. Only the e-value is considered during this stage. Ortholog clusters where e-values meet a user-defined threshold to sequences in the ORAdb proceed to the restrictive search. The default e-value was set to 0.001, as the main objective of the relaxed search is to capture as many sequences for annotation as possible while avoiding an exaggerated number of sequence alignments. In the restrictive search, all sequences in the transitioned ortholog clusters are aligned to all the sequences in the reaction set(s) present in the ORAdb to which they had a hit during the relaxed search. Again, the query sequence is only assigned to the function of a reference sequence if the e-value is below a determined threshold (default 1e^-9^). Next, an additional filter is applied based on annotation bit score values (default 50). Although we established default values for the relaxed and restrictive search as well as bit score, the user has the option to define the thresholds for all individual parameters.

The identification of putative interactions between species is based on all combinations of bacterial isolates with the genomic content to perform the user-defined pathway defined in the ORAdb. The input for this task consists of: (1) a binary table generated at the end of the functional annotation which indicates the presence or absence of sequences annotated to each reaction in the ORAdb in each species (e.g. Supplementary Table S10); (2) a set of Gene-Protein-Reaction (GPR) rules for all reactions considered (e.g. Supplementary data - Table S20); and (3) a user-defined tab-delimited file where the sets of reactions for complete pathways, subsets of reactions required to be performed by single species and transporter-associated genes (e.g. Supplementary data – Table S1) are described. To further reduce the vast amount of putative microbial interactions and to increase confidence in the results manual filtering can be performed to reflect available knowledge (e.g. known cross-feeding relationship between species) and/or the likelihood of biologically feasible species interactions). The user also may have interest in assessing subsets of microbial interactions using specific criteria. Therefore, additional constraints can be applied to the list of putative microbial interactions further reducing the search space. These include the degree of completeness of a pathway, the number of reactions expected to be performed by a single species or the presence or absence of transporter genes. Additionally, a graphical network visualization is also produced during this step. Graphical network visualization is implemented in R using the packages visNetwork (v2.0.9), reshape2 (v1.4.3) and RColorBrewers (v1.1-2) but also requires the pandoc linux library. Graphical visualization was implemented with R v3.6 but tested also with v4.0. The visualization creates a HTML file that allows interactive exploration of the network and provides hyperlinks to KEGG if available.

All tasks - functional annotation, prediction of putative microbial interactions and generation of graphical visualizations - are performed by running the script *annotate_and_predict*.*sh* included in OrtSuite (https://github.com/mdsufz/OrtSuite/blob/master/annotate_and_predict.sh). OrtSuite’s predictions of individual species and combinations of species with the genetic potential to perform each defined pathway is stored in text files located in a folder termed “interactions”.

### Conversion of benzoate to acetyl-CoA as a model pathway

We selected three alternative pathways involved in the conversion of benzoate to acetyl-CoA (BTA) to test the functional annotation and prediction of putative synergistic microbial interactions using OrtSuite (Supplementary data - Table S14). Two pathways consisted in the anaerobic degradation of benzoate to acetyl-CoA via benzoyl-CoA differing only in the reactions required for transformation of glutaryl-CoA to crotonyl-CoA (hereafter, respectively, P1 and P2). P1 first converts glutaryl-CoA to glutaconyl-CoA and then to crotonoyl-CoA while P2 directly converts glutaryl-CoA to crotonoyl-CoA. One pathway consisted in the aerobic degradation of benzoate via catechol (hereafter P3). The complete number of reactions, enzymes, KO identifiers and KO-associated sequences in each alternative pathway is shown in the supplementary data (Supplementary data - Table S25).

### Species selection for testing functional annotation

To assess the performance of OrtSuite, we selected the transformation of benzoate to acetyl-CoA as a model pathway and a set of previously characterized species known to be involved in this pathway (Table 1). This set of species was divided in two groups. The first group contained sequenced genomes of species whose ability to convert benzoate to acetyl-CoA has been demonstrated by KEGG (Kanehisa *et al*, 2004) and were selected as positive controls. These species were classified according to their genomic potential: complete, if all protein encoding genes required for a BTA pathway were present in their genome or partial, if not all protein encoding genes were present. The second group consisted of species known to lack all required protein encoding genes and were selected as negative controls. In total, we selected 18 species as positive controls. Seven of them have the genetic potential to perform the alternative P2 pathway; eight have the genetic potential to perform alternative path P3 (positive controls); and, none able to completely perform the alternative path P1. To note that species *Thauera sp*. MZ1T has the genetic potential to perform P2 and P3 pathways. Four organisms were selected as negative controls. Using their genomes, we evaluated the performance of OrtSuite based on precision and recall rates for clustering of orthologs and the correct functional annotation of sequences. Also, a set of genomes from the species containing the genetic potential to degrade benzoate were artificially mutated at the nucleotide level at different rates in order to determine how levels of point mutations in open reading frames (ORFs) affected clustering of ortholog groups.

### Species selection for validation of putative interspecies interactions

In a study performed by Fetzer and collaborators (Fetzer *et al*, 2015) community biomass production of mono- and mixed-cultures was assessed in medium containing benzoate. The authors used this data to infer potential species interactions. This set of genomes was processed with OrtSuite to determine the species’ genetic potential to degrade benzoate, either individually or as a result of their interaction. Our results were compared to those obtained by Fetzer and collaborators and used to assess whether the study’s inferred potential interactions could be derived from their combined genetic potential.

### Evaluation of ortholog clustering

The clustering of orthologs was evaluated by measuring the pairwise precision and recall. Clustering precision measures how many pairs of sequences associated with the same molecular function are grouped together and is calculated by dividing the number of correctly clustered sequences by the total number of clustered sequences (Equation 1).

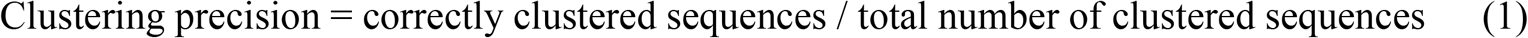

where, correctly clustered sequences refers to the pairs of sequences that share the same function and are clustered together and total number of clustered sequences refers to all pairs of sequences that are clustered together irrespective of sharing the same function.

Clustering recall measures how many pairs of sequences with the same molecular function are not clustered together. Recall is calculated by dividing the number of correctly clustered sequences by the total true sequence clusters (Equation 2).

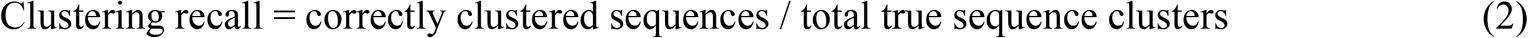

where, correctly clustered sequences refers to the pairs of sequences that share the same function and are clustered together and total true sequence clusters refers to all pairs of sequences that have the same function.

### Evaluation of sequence aligner used for clustering of orthologs

Changes of a single DNA base can result in the production of a different amino acid which might result in a different protein. To determine the impact of mutations on the clustering of orthologs a single gene from three species was artificially mutated at different rates. These mutations were introduced in the nucleotide sequences of each gene. Only substitutions were considered since these are the most commonly studied (Lynch, 2010) and none of the mutations were allowed to occur on the first and last codon. When, during the mutation, new stop or/and start codons were introduced, the translation was made for all the possible proteins and the largest was selected.

*Burkholderia vietnamiensis* G4 was mutated on the gene K05783, *Azoarcus sp*. CIB on the gene K07537 and *Aromatoleum aromaticum* EbN1 on the gene K07538. Each gene was mutated at rates of 0.01, 0.03, 0.05, 0.1, 0.15 and 0.25. Each mutation rate resulted in an in silico strain of the original genome (e.g., *Burkholderia vietnamiensis* G4 strain K05783_25, where “K05783” is the KEGG ortholog identifier and “25” is the rate of mutation). A total of 18 strains were generated (six in silico mutated strains per genome). The complete set of original and artificially mutated genomes is available in a compressed file (Supplementary data - Test_genomes_set.zip).

### Evaluation of functional annotation

Functional annotation was evaluated based on the data collected from KEGG (Altschul *et al*, 1990). Annotation performance is calculated by dividing the number of matching annotated sequences by the total number of annotations (Equation 3).

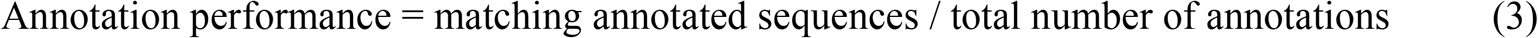

where, matching annotated sequences refers to the number of sequences annotated by KEGG annotations predicted by OrtSuite and total number of annotations refers to the all sequences that were assigned a function by KEGG.

### Evaluation of microbial interaction predictions

We evaluated the prediction of putative microbial interactions using a genome set from an independent study (Fetzer *et al*, 2015) containing species with exhibited growth in medium containing benzoate (defined as Fetzer_genome_set). The authors do not identify specific potential interactions in the transformation of benzoate but infer interspecific interactions in an environment containing benzoate as the major carbon source. For the complete set of species combinations and benzoate degradation capabilities and effects identified by Fetzer and collaborators, see (Fetzer *et al*, 2015) (Supplementary data - Table S24).

#### Bacterial cultures and sequencing

Bacterial cryo-cultures of the different isolates were revived on LB agar plates. Single colonies were picked and grown overnight in 2 ml LB medium at 37°C. The cells were pelleted by centrifugation. Cells were lysed and genomic DNA was extracted using a Nucleospin Tissue Kit (Machery and Nagel). Approximately 150 to 1000 ng of DNA were used for fragmentation (insert size: 300 – 700 bp) and sequencing libraries were prepared following the NEB Ultra II FS Kit protocol (New England Biolabs). Libraries were quantified using a JetSeq Library Quantification Lo-ROX Kit (Bioline) and quality-checked by Bioanalyzer (Agilent). These libraries were sequenced on an Illumina MiSeq instrument with a final concentration of 8 pM using the v3 600 cycles chemistry and 5% PhiX.

#### Genome assembly and Open Reading Frame prediction

The sequenced reads were quality checked using Trim Galore v0.4.4_dev. Next, genomes were assembled using the Spades Assembler v3.15.2 and their quality assessed using CheckM. Taxonomic classification was performed using Genome Taxonomy Database (GTDBTk) release Open Reading Frames (ORFs) were predicted using Prodigal v2.6.3. Translation of sequences to amino acid format was performed using faTrans from kentUtils (https://github.com/ENCODE-DCC/kentUtils/tree/master/src/utils/faTrans).

## Author Contributions

JS, OD, PS and UNR developed the concept of OrtSuite. JS, MG, AB and UNR developed the OrtSuite workflow. JS, MG, AB and UNR performed the benchmarks. CV provided information and data for defining benzoate to acetyl-CoA conversion pathways. RK sequenced bacterial isolates that where provided by AC. AB created the interactive network visualization module. JS and UNR wrote the manuscript. All authors read and commented on different versions of the manuscript and approved the final manuscript.

## Data Availability

The datasets and computer code produced in this study are available in the following databases:

- The genomes used to test the workflow are available at National Centre for Biotechnology Information (https://www.ncbi.nlm.nih.gov/) under the accession identifiers CP029389-CP029397, GCF_000001735, AP012304, AP012305, CP021731, CP011072, CP007785-CP007787, CP000614-CP000621, CP003230, CP005996, CP003108, CR555306-CR5553068, GCF_000225785, LN997848-LN997849, CP022989-CP022996, CP024315, AP012547, CP022046-CP022047 and CP001281-CP001282.
- The genome assemblies used to predict interspecies interactions are available at National Centre for Biotechnology Information (https://www.ncbi.nlm.nih.gov/) with the study accession PRJEB38476: (https://www.ncbi.nlm.nih.gov/bioproject/648592).
- OrtSuite scripts: GitHub (https://github.com/mdsufz/OrtSuite).

## Acknowledgments

We thank the early users of OrtSuite Sandra Silva, Felipe Côrrea and Jonas Kasmanas for their help with debugging and for workflow suggestions. We also thank Diogo Lima and Emanuel Cunha for their assistance in the implementation of the script required to generate the Gene-Protein-Reaction (GPR) rules; and Nicole Steinbach for her work in the sequencing of the isolates used as the test set.

## References

Altschul SF, Gish W, Miller W, Myers EW & Lipman DJ (1990) Basic local alignment search tool. Journal of molecular biology 215: 403–10

Bazire A, Diab F, Jebbar M & Haras D (2007) Influence of high salinity on biofilm formation and benzoate assimilation by Pseudomonas aeruginosa. Journal of Industrial Microbiology and Biotechnology 34: 5–8

Buchfink B, Xie C & Huson DH (2015) Fast and sensitive protein alignment using DIAMOND. Nat Methods 12: 59–60

Delmas E, Besson M, Brice M-H, Burkle LA, Riva GVD, Fortin M-J, Gravel D, Guimarães PR, Hembry DH, Newman EA, et al (2019) Analysing ecological networks of species interactions. Biological Reviews 94: 16–36

Devanadera A, Vejarano F, Zhai Y, Suzuki-Minakuchi C, Ohtsubo Y, Tsuda M, Kasai Y, Takahata Y, Okada K & Nojiri H (2019) Complete Genome Sequence of an Anaerobic Benzene-Degrading Bacterium, Azoarcus sp. Strain DN11. Microbiol Resour Announc 8

Dong X & Strous M (2019) An Integrated Pipeline for Annotation and Visualization of Metagenomic Contigs. Front Genet 10

Emms DM & Kelly S (2015) OrthoFinder: solving fundamental biases in whole genome comparisons dramatically improves orthogroup inference accuracy. Genome biology 16: 157–157

Emms DM & Kelly S (2019) OrthoFinder: phylogenetic orthology inference for comparative genomics. Genome Biology 20: 238

Fetzer I, Johst K, Schäwe R, Banitz T, Harms H & Chatzinotas A (2015) The extent of functional redundancy changes as species’ roles shift in different environments. Proc Natl Acad Sci USA 112: 14888–14893

Gabaldón T & Koonin EV (2013) Functional and evolutionary implications of gene orthology. Nature Reviews Genetics 14: 360–366

Gottstein W, Olivier BG, Bruggeman FJ & Teusink B (2016) Constraint-based stoichiometric modelling from single organisms to microbial communities. Journal of The Royal Society Interface 13: 20160627

Heirendt L, Arreckx S, Pfau T, Mendoza SN, Richelle A, Heinken A, Haraldsdóttir HS, Wachowiak J, Keating SM, Vlasov V, et al (2019) Creation and analysis of biochemical constraint-based models using the COBRA Toolbox v.3.0. Nature Protocols 14: 639–702

Hernández-Salmerón JE & Moreno-Hagelsieb G (2020) Progress in quickly finding orthologs as reciprocal best hits: comparing blast, last, diamond and MMseqs2. BMC Genomics 21: 741

Hu Y, Feng Y, Zhang X & Zong Z (2017) Acinetobacter defluvii sp. nov., recovered from hospital sewage. International Journal of Systematic and Evolutionary Microbiology, 67: 1709–1713

Huerta-Cepas J, Forslund K, Coelho LP, Szklarczyk D, Jensen LJ, von Mering C & Bork P (2017) Fast Genome-Wide Functional Annotation through Orthology Assignment by eggNOG-Mapper. Mol Biol Evol 34: 2115–2122

Jenul C, Sieber S, Daeppen C, Mathew A, Lardi M, Pessi G, Hoepfner D, Neuburger M, Linden A, Gademann K, et al (2018) Biosynthesis of fragin is controlled by a novel quorum sensing signal. Nat Commun 9: 1–13

Junghare M, Patil Y & Schink B (2015) Draft genome sequence of a nitrate-reducing, o-phthalate degrading bacterium, Azoarcus sp. strain PA01T. Standards in Genomic Sciences 10: 90

Kanehisa M, Goto S, Kawashima S, Okuno Y & Hattori M (2004) The KEGG resource for deciphering the genome. Nucleic acids research 32: D277–D280

Kanehisa M, Sato Y & Morishima K (2016) BlastKOALA and GhostKOALA: KEGG Tools for Functional Characterization of Genome and Metagenome Sequences. J Mol Biol 428: 726–731

Khanal A, Yu McLoughlin S, Kershner JP & Copley SD (2015) Differential Effects of a Mutation on the Normal and Promiscuous Activities of Orthologs: Implications for Natural and Directed Evolution. Mol Biol Evol 32: 100–108

Kitko RD, Cleeton RL, Armentrout EI, Lee GE, Noguchi K, Berkmen MB, Jones BD & Slonczewski JL (2009) Cytoplasmic Acidification and the Benzoate Transcriptome in Bacillus subtilis. PLOS ONE 4: e8255

Koonin EV (2005) Orthologs, Paralogs, and Evolutionary Genomics. Annual Review of Genetics 39: 309– 338

Lee Y, Lee Y & Jeon CO (2019) Biodegradation of naphthalene, BTEX, and aliphatic hydrocarbons by Paraburkholderia aromaticivorans BN5 isolated from petroleum-contaminated soil. Sci Rep 9: 860

Li L, Stoeckert CJ & Roos DS (2003) OrthoMCL: identification of ortholog groups for eukaryotic genomes. Genome Res 13: 2178–2189

Li W & Godzik A (2006) Cd-hit: a fast program for clustering and comparing large sets of protein or nucleotide sequences. Bioinformatics 22: 1658–1659

Locey KJ & Lennon JT (2016) Scaling laws predict global microbial diversity. PNAS: 201521291

Lynch M (2010) Evolution of the mutation rate. Trends Genet 26: 345–352

Lyu Z, Shao N, Akinyemi T & Whitman WB (2018) Methanogenesis. Curr Biol 28: R727–R732

Madden T (2003) The BLAST Sequence Analysis Tool National Center for Biotechnology Information (US)

Maestre FT, Castillo-Monroy AP, Bowker MA & Ochoa-Hueso R (2012) Species richness effects on ecosystem multifunctionality depend on evenness, composition and spatial pattern. Journal of Ecology 100: 317–330

Mendes LW, Raaijmakers JM, de Hollander M, Mendes R & Tsai SM (2018) Influence of resistance breeding in common bean on rhizosphere microbiome composition and function. ISME J 12: 212– 224

Messina E, Denaro R, Crisafi F, Smedile F, Cappello S, Genovese M, Genovese L, Giuliano L, Russo D, Ferrer M, et al (2016) Genome sequence of obligate marine polycyclic aromatic hydrocarbons-degrading bacterium Cycloclasticus sp. 78-ME, isolated from petroleum deposits of the sunken tanker Amoco Milford Haven, Mediterranean Sea. Marine Genomics 25: 11–13

Meyer-Cifuentes I, Fiedler S, Müller JA, Kappelmeyer U, Mäusezahl I & Heipieper HJ (2017) Draft Genome Sequence of Magnetospirillum sp. Strain 15-1, a Denitrifying Toluene Degrader Isolated from a Planted Fixed-Bed Reactor. Genome Announc 5

Mistry J, Chuguransky S, Williams L, Qureshi M, Salazar GA, Sonnhammer ELL, Tosatto SCE, Paladin L, Raj S, Richardson LJ, et al (2021) Pfam: The protein families database in 2021. Nucleic Acids Research 49: D412–D419

Morin M, Pierce EC & Dutton RJ (2018) Changes in the genetic requirements for microbial interactions with increasing community complexity. eLife 7: e37072

Mrozik A & Labuzek S (2002) A comparison of biodegradation of phenol and homologous compounds by Pseudomonas vesicularis and Staphylococcus sciuri strains. Acta Microbiol Pol 51: 367–378

Mulder CPH, Uliassi DD & Doak DF (2001) Physical stress and diversity-productivity relationships: The role of positive interactions. PNAS 98: 6704–6708

Ng PC & Henikoff S (2006) Predicting the effects of amino acid substitutions on protein function. Annu Rev Genomics Hum Genet 7: 61–80

O’Sullivan LA, Weightman AJ, Jones TH, Marchbank AM, Tiedje JM & Mahenthiralingam E (2007) Identifying the genetic basis of ecologically and biotechnologically useful functions of the bacterium Burkholderia vietnamiensis. Environmental Microbiology 9: 1017–1034

Parks DH, Rinke C, Chuvochina M, Chaumeil P-A, Woodcroft BJ, Evans PN, Hugenholtz P & Tyson GW (2017) Recovery of nearly 8,000 metagenome-assembled genomes substantially expands the tree of life. Nat Microbiol 2: 1533–1542

Pasolli E, Asnicar F, Manara S, Zolfo M, Karcher N, Armanini F, Beghini F, Manghi P, Tett A, Ghensi P, et al (2019) Extensive Unexplored Human Microbiome Diversity Revealed by Over 150,000 Genomes from Metagenomes Spanning Age, Geography, and Lifestyle. Cell 176: 649-662.e20

Pearson WR (2013) An Introduction to Sequence Similarity (“Homology”) Searching. Curr Protoc Bioinformatics 0 3

Peng T, Luo A, Kan J, Liang L, Huang T & Hu Z (2018) Identification of A Ring-Hydroxylating Dioxygenases Capable of Anthracene and Benz[a]anthracene Oxidization from Rhodococcus sp. P14. MMB 28: 183–189

Prestat E, David MM, Hultman J, Taş N, Lamendella R, Dvornik J, Mackelprang R, Myrold DD, Jumpponen A, Tringe SG, et al (2014) FOAM (Functional Ontology Assignments for Metagenomes): a Hidden Markov Model (HMM) database with environmental focus. Nucleic Acids Res 42: e145

Rabus R, Boll M, Heider J, Meckenstock RU, Buckel W, Einsle O, Ermler U, Golding BT, Gunsalus RP, Kroneck PMH, et al (2016) Anaerobic Microbial Degradation of Hydrocarbons: From Enzymatic Reactions to the Environment. MMB 26: 5–28

Ravikrishnan A & Raman K (2021) Unraveling microbial interactions in the gut microbiome. bioRxiv: 2021.05.17.444446

Raynaud X & Nunan N (2014) Spatial Ecology of Bacteria at the Microscale in Soil. PLOS ONE 9: e87217

Robertson WJ, Franzmann PD & Mee BJ (2000) Spore-forming, Desulfosporosinus-like sulphate-reducing bacteria from a shallow aquifer contaminated with gasolene. Journal of Applied Microbiology 88: 248–259

Roh SW, Abell GCJ, Kim K-H, Nam Y-D & Bae J-W (2010) Comparing microarrays and next-generation sequencing technologies for microbial ecology research. Trends Biotechnol 28: 291–299

Seemann T (2014) Prokka: rapid prokaryotic genome annotation. Bioinformatics 30: 2068–2069

Singleton CM, Petriglieri F, Kristensen JM, Kirkegaard RH, Michaelsen TY, Andersen MH, Kondrotaite Z, Karst SM, Dueholm MS, Nielsen PH, et al (2021) Connecting structure to function with the recovery of over 1000 high-quality metagenome-assembled genomes from activated sludge using long-read sequencing. Nat Commun 12: 2009

Slade EM, Kirwan L, Bell T, Philipson CD, Lewis OT & Roslin T (2017) The importance of species identity and interactions for multifunctionality depends on how ecosystem functions are valued. Ecology 98: 2626–2639

Sperfeld M, Diekert G & Studenik S (2019) Anaerobic aromatic compound degradation in Sulfuritalea hydrogenivorans sk43H. FEMS Microbiol Ecol 95

Suvorova IA & Gelfand MS (2019) Comparative Genomic Analysis of the Regulation of Aromatic Metabolism in Betaproteobacteria. Front Microbiol 10

Tal O, Selvaraj G, Medina S, Ofaim S & Freilich S (2020) NetMet: A Network-Based Tool for Predicting Metabolic Capacities of Microbial Species and their Interactions. Microorganisms 8: 840

Thommes M, Wang T, Zhao Q, Paschalidis IC & Segrè D (2019) Designing Metabolic Division of Labor in Microbial Communities. mSystems 4

Tully BJ, Graham ED & Heidelberg JF (2018) The reconstruction of 2,631 draft metagenome-assembled genomes from the global oceans. Scientific Data 5: 170203

Valderrama JA, Durante-Rodríguez G, Blázquez B, García JL, Carmona M & Díaz E (2012) Bacterial Degradation of Benzoate: CROSS-REGULATION BETWEEN AEROBIC AND ANAEROBIC PATHWAYS. J Biol Chem 287: 10494–10508

Wang B, Lai Q, Cui Z, Tan T & Shao Z (2008) A pyrene-degrading consortium from deep-sea sediment of the West Pacific and its key member Cycloclasticus sp. P1. Environmental Microbiology

Wang W, Yang S, Hunsinger GB, Pienkos PT & Johnson DK (2014) Connecting lignin-degradation pathway with pre-treatment inhibitor sensitivity of Cupriavidus necator. Frontiers in Microbiology 5: 247

